## Supplementary material for "The neurocognitive role of working memory load when Pavlovian motivational control affects instrumental learning": TextS2.docx

**The effect of WM load on Pavlovian bias (hypothesis 2)**

We found no main effect of Pavlovian bias between the GNG and WMGNG task (WMGNG > GNG [Pavlovian congruent > Pavlovian incongruent]) within the striatum or the SN/VTA (*p* < 0.05 SVC). There was no significant voxel within SN/VTA whose *p* > 0.001 (uncorrected), thus here we report neural statistics within the striatum only (**Table S5**).

**The effect of WM load on choice randomness (hypothesis 3)**

We observed no main effect of WM load on choice randomness (WMGNG > GNG [W_chosen_ - W_unchosen_]) within the vmPFC (*p* < 0.05 SVC). There was no significant voxel within vmPFC whose *p* > 0.001 (uncorrected).

**Replication analyses**

We attempted to replicate findings in the literature, focusing on the main effect of WM load, reward, and loss (**Fig S6**, **Table S3**) to validate our approach. We used the GLM2 model and then three contrasts were computed for replication: the main effect of WM load ([(1) + (2) + (3) + (4)]_WMGNG_ - [(1)+(2)+(3)+(4)]_GNG_), the main effect of reward (irrespective of the type of task, (7) - (8)), the main effect of loss (irrespective of the type of task, (9) - (8)).
