## Supplementary figures and images for "The neurocognitive role of working memory load when Pavlovian motivational control affects instrumental learning"

### FigureS1.png

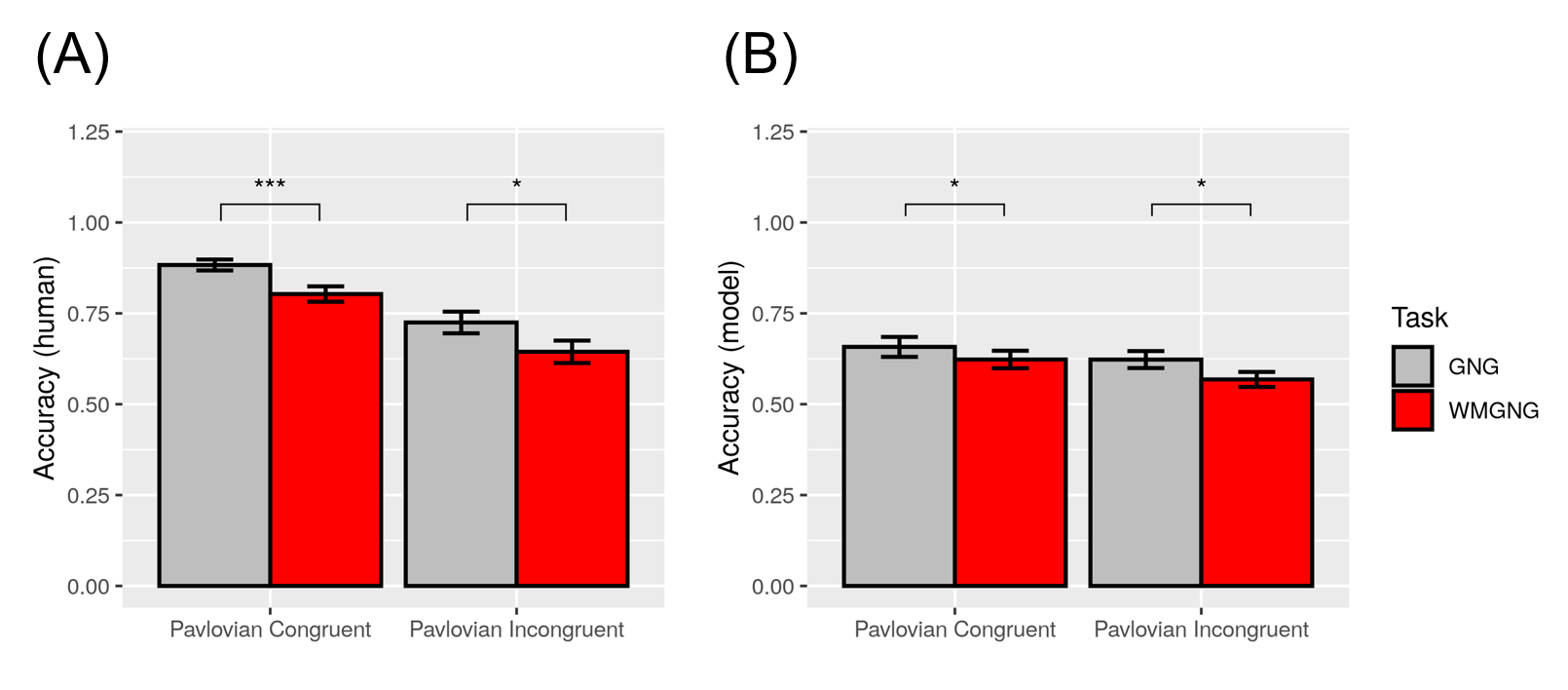

### FigureS2.png

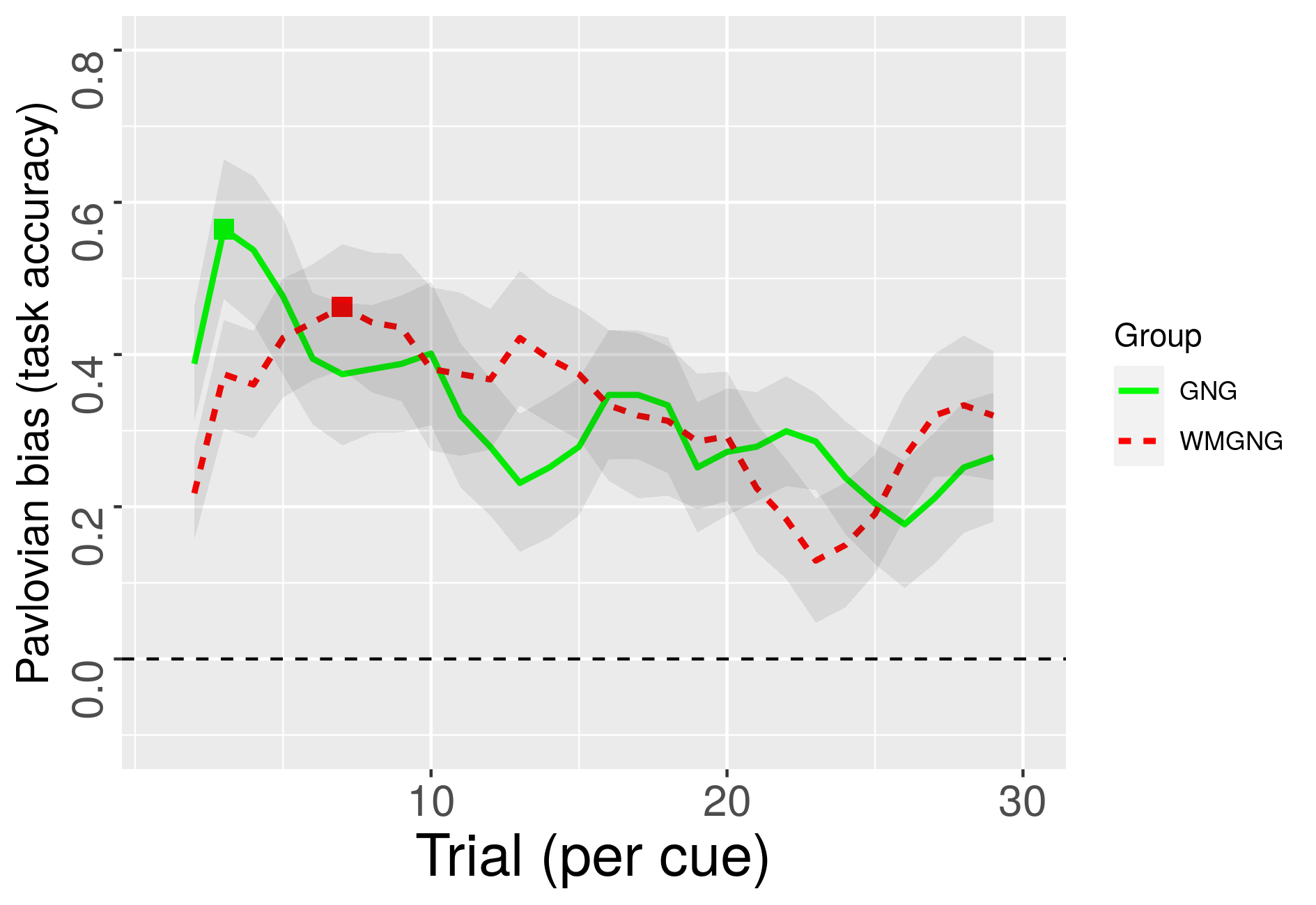

### FigureS3.png

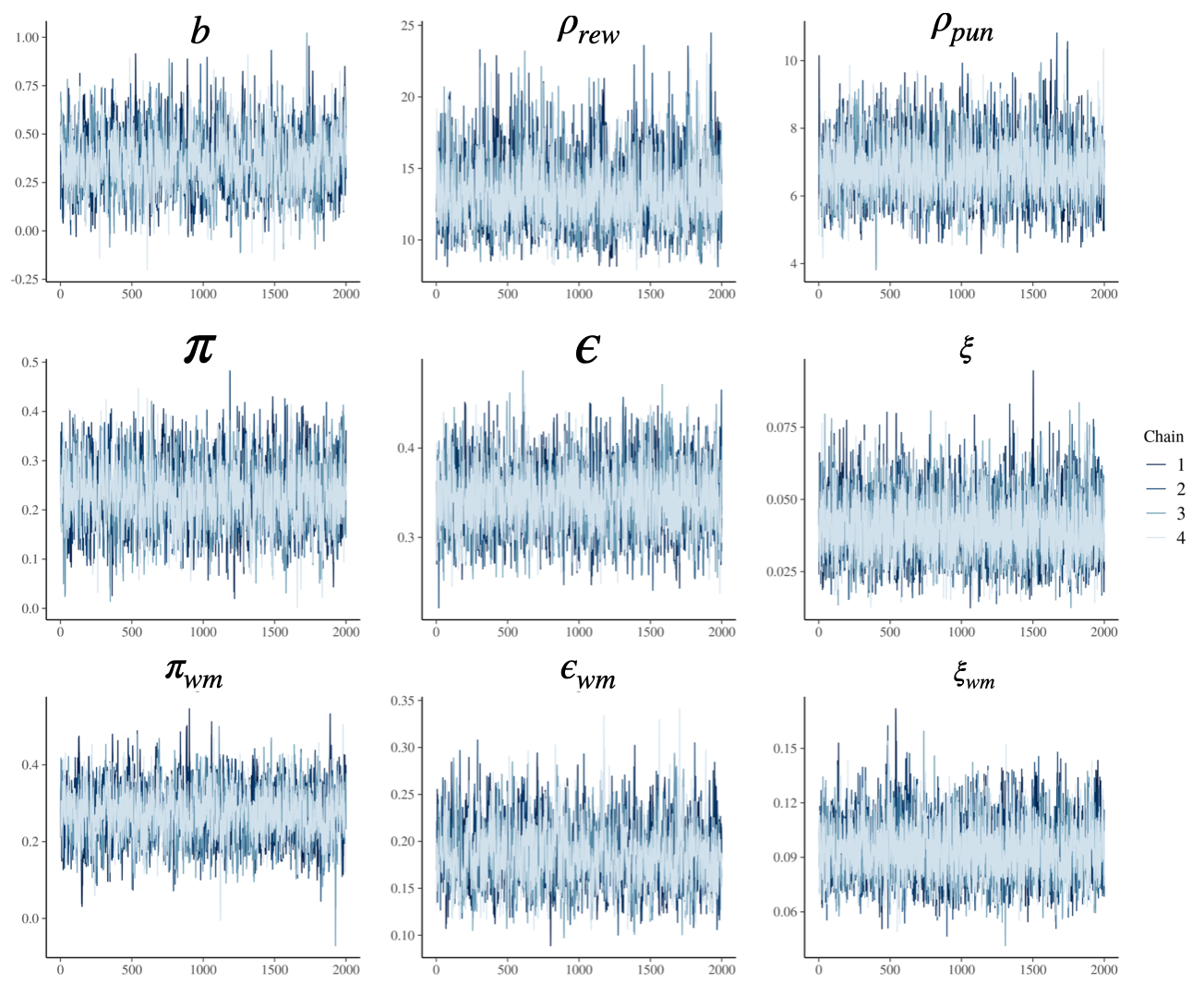

### FigureS4.png

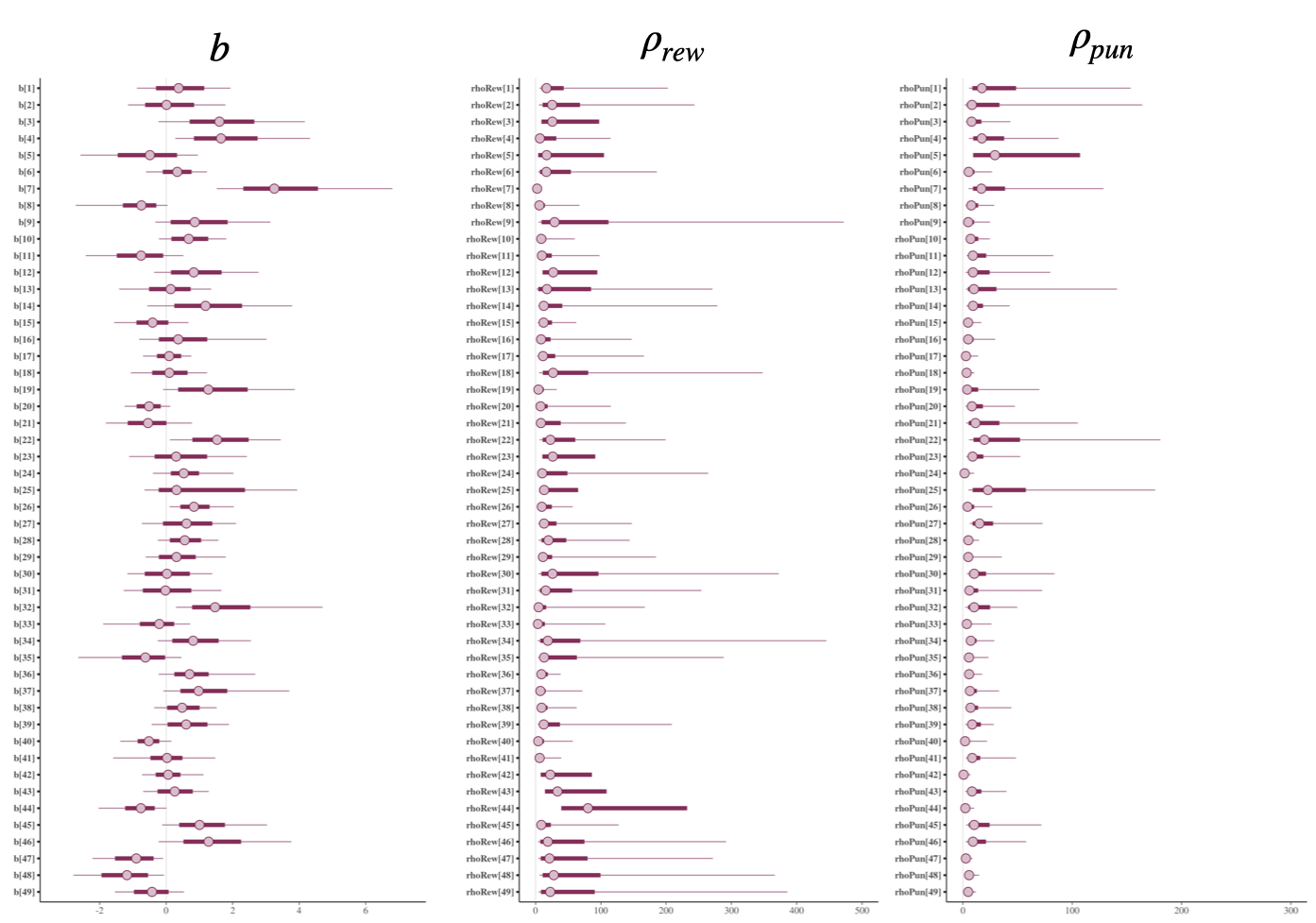

### FigureS5.png

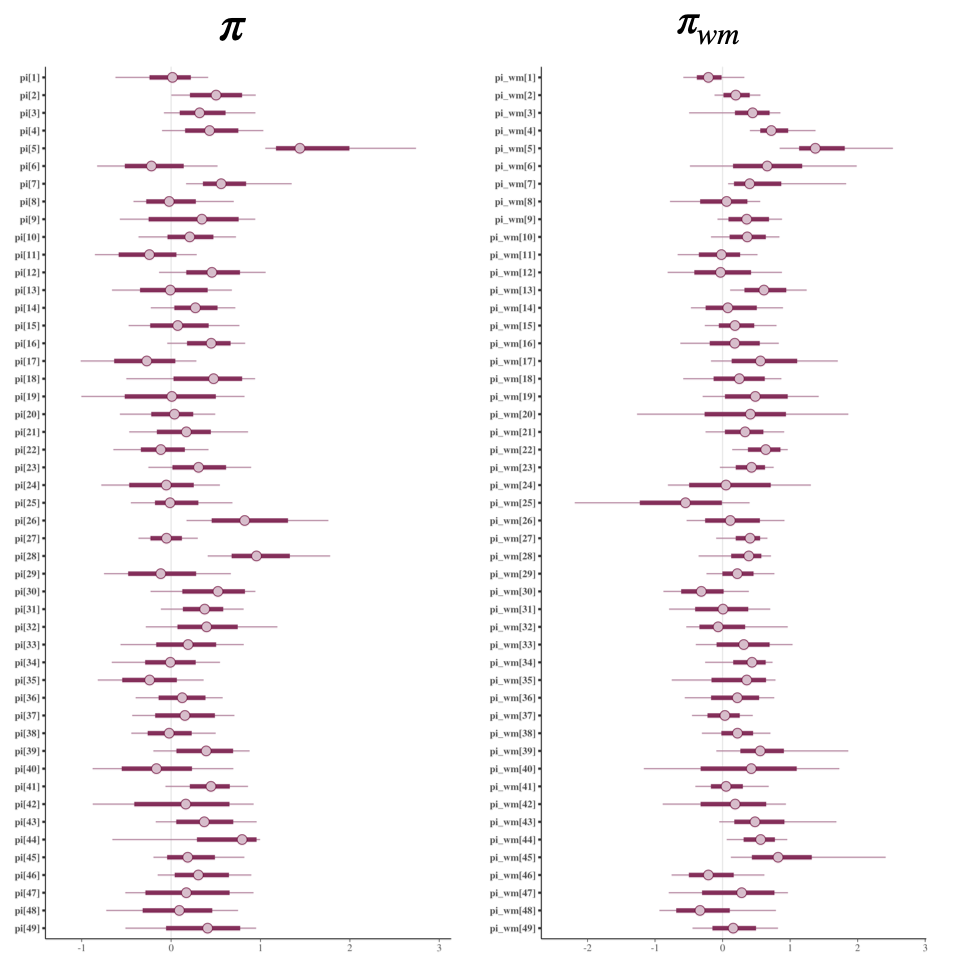

### FigureS6.png

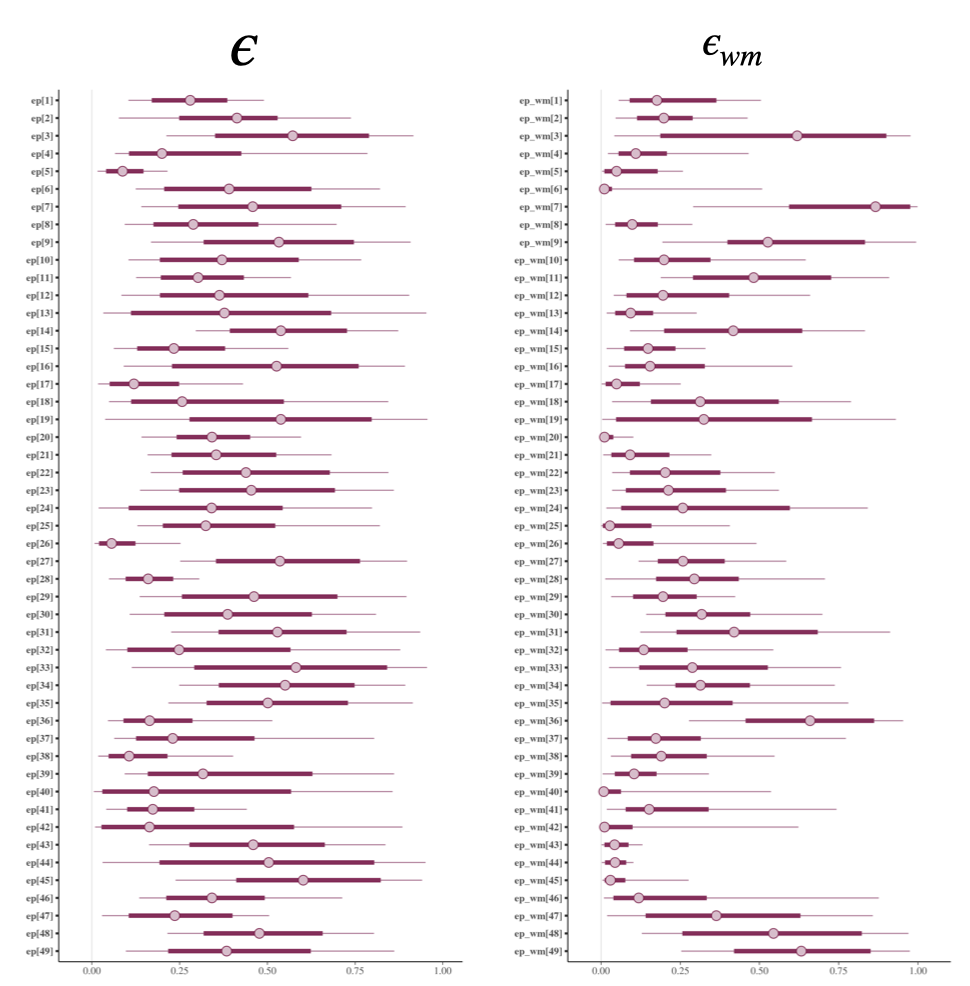

### FigureS7.png

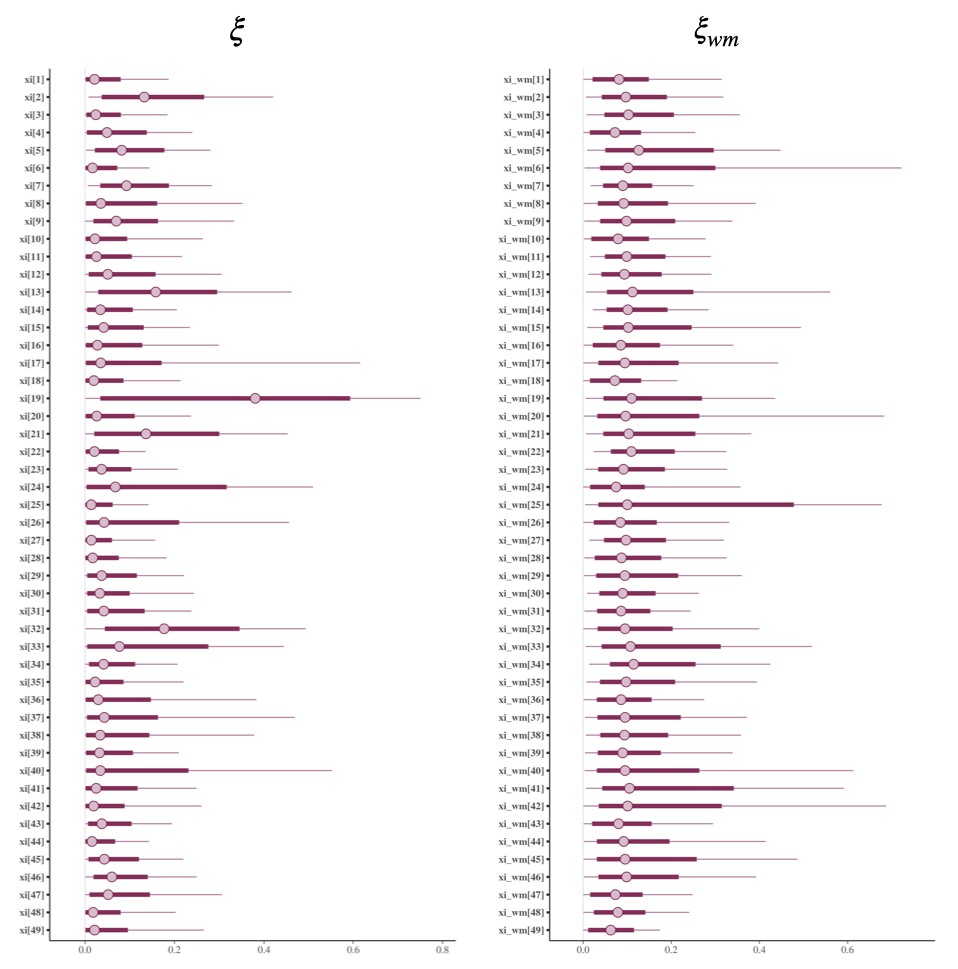

### FigureS8.png

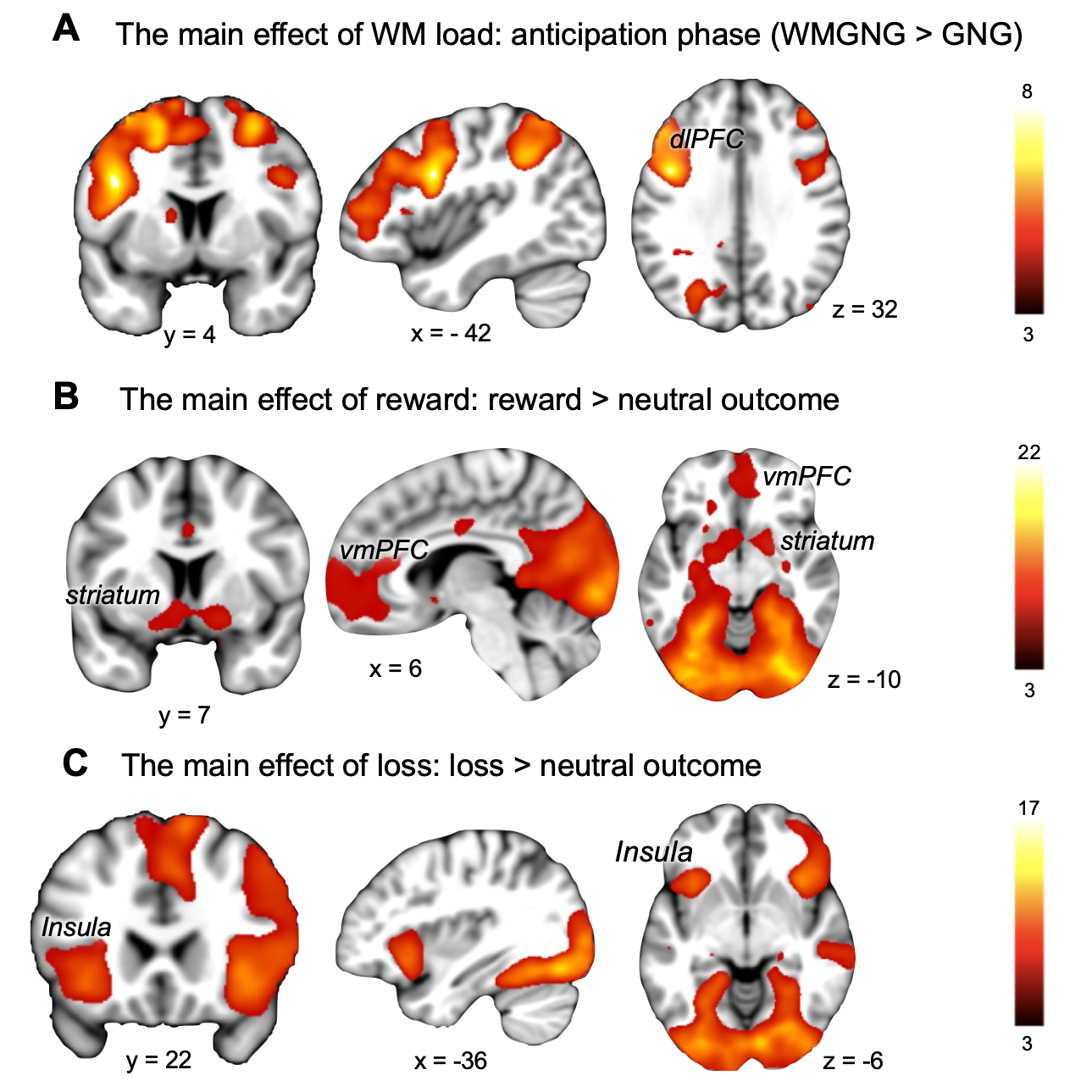
